# Seasonal resistome diversity and dissemination of WHO priority antibiotic-resistant pathogens in Lebanese estuaries

**DOI:** 10.1101/2021.12.13.472353

**Authors:** Wadad Hobeika, Margaux Gaschet, Marie-Cécile Ploy, Elena Buelow, Dolla Karam Sarkis, Christophe Dagot

## Abstract

Anthropogenic activities are demonstrated to be the key drivers of antimicrobial resistance (AMR) dissemination in the environment. Lebanese rivers that lead to the Mediterranean Sea were sampled at estuaries’ sites, under high anthropogenic pressure, in spring 2017 and winter 2018 to study seasonal variation of antimicrobial-resistant bacteria (ARBs) and antimicrobial resistance genes (ARGs). Methods: A combined approach using culture techniques and high throughput qPCR identified hotspots for antimicrobial resistance and anthropogenic pressure in particular locations along the Lebanese coast. Results: Multi-resistant Gram-negative (Enterobacterales and *Pseudomonas* spp) and Gram-positive bacterial pathogens were isolated. A high abundance of certain ARGs (*vanB, bla*_BIC-1_, *bla*_GES_, tetM, and *mcr-1)* was detected in 5 Lebanese estuaries. The relative abundance of ARGs was highest in winter and areas with high anthropogenic activities and population growth with an influx of refugees. Conclusion: Qualitative analysis of ARB and the analysis of the Lebanese estuaries’ resistome revealed critical levels of contamination with pathogenic bacteria and provided significant information about the spread of ARGs in anthropogenically impacted estuaries.

## Introduction

Antimicrobial resistance (AMR) is a One Health crisis aggravated by the lack of water and pollution management on a global scale ^1,2^.Anthropogenic activities are demonstrated to be the key drivers of AMR dissemination in the environment ^3,4,5,6^ subsequently altering its ecosystems ^7,8,9^. Through discharge of treated or untreated wastewater (WW) effluents into surface water, a high abundance of antibiotic resistance genes (ARGs), antibiotic-resistant bacteria (ARBs), and mobile genetic elements (MGEs) mixed with a cocktail of micropollutants and drug residues are continuously disseminated into the environment ^10,11,12,13^. Additionally, agricultural practices such as soil fertilization with manure and sludge, or irrigation with WW effluents, further expand the environmental background levels of pollutants associated with the dissemination of antimicrobial resistance ^14,9^. In Lebanon, rules to control the overuse and misuse of antibiotics for treatment, growth promotion, and prophylaxis in agriculture and animal husbandry are not strictly implemented ^15,16^ which would likely contribute to increasing AMR in the Lebanese and connected environments ^17,18^.

The surveillance of AMR in the environment is often assessed through ARG quantification ^19,20^. High abundances of ARGs were found to be associated with fecal contamination ^21,22,23^.

Estuaries are transitional zones between rivers and sea bodies ^24,25^ exhibiting properties of marine and freshwater and underlining continental-oceanic interactions ^26^. Estuaries are also considered filtering and buffered ecosystems broadly anthropized ^24,27^. Contaminants can reside for prolonged periods due to the tidal streams ^27,28^. The spatial-temporal dissemination of ARB and ARGs in estuaries remains understudied ^26^. The Mediterranean coastline is densely populated with a large anthropic footprint, i.e. intensive fishing, shipping, recreational and industrial activities ^29^. Our study aims to identify the impact of anthropogenic activities in AMR dissemination in the Mediterranean Sea through the Lebanese estuaries. The objectives of this study were to monitor in the Lebanese river estuaries, the dissemination of i) ARBs, notably extended-spectrum β-lactamase-producing-Enterobacterales (ESBL-E), multi-drug resistant *Pseudomonas aeruginosa*, and methicillin-resistant *Staphylococcus aureus* (MRSA), and ii) the resistome through the detection of ARGs and MGEs.

## Results

### Bacterial culture

In total, we isolated 19 different bacterial species in different quantities in spring and winter from the estuarine water samples along the Lebanon coastline (see the Materials and Methods section) (Table 1). From 1 mL of the water samples cultivated on selective media, we obtained 50 and 10064 CFUs (colony forming units) resistant Enterobacterales and 41 and 43 CFUs resistant *Pseudomonas spp*. in spring and winter respectively. However, among the 10064 CFUs in winter, 10^4^ corresponded to the same bacterial species (*Hafnia alvei*) in the Beirut estuary. A more diverse panel of Enterobacterales species was isolated in spring. Enterobacetrales and/or *Pseudomonas spp*. strains were isolated from all samples, except the Bared river. Moreover, Enterobacterales were not detected in the Kaleb river and *Pseudomonas* spp were not found in the Zahnari river. MRSA isolates were detected only in spring.

**Table 1.**
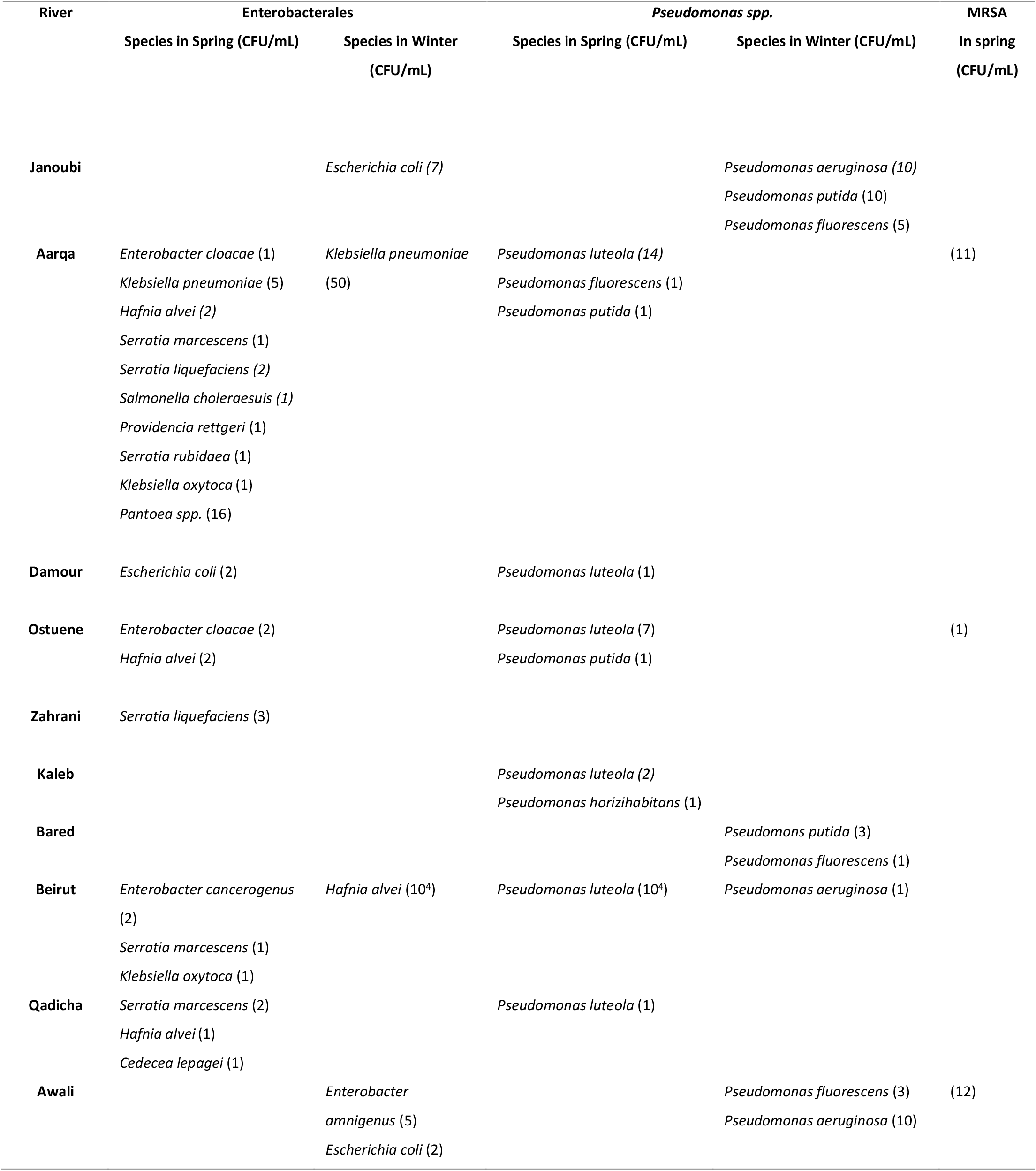
Gram-negative bacterial species isolated in the rivers’ estuaries

| River | Enterobacterales |  | Pseudomonas spp. |  | MRSA |
| --- | --- | --- | --- | --- | --- |
|  | Species in Spring (CFU/mL) | Species in Winter (CFU/mL) | Species in Spring (CFU/mL) | Species in Winter (CFU/mL) | In spring (CFU/mL) |
| Janoubi |  | *Escherichia coli* (7) |  | *Pseudomonas aeruginosa* (10)<br>*Pseudomonas putida* (10)<br>*Pseudomonas fluorescens* (5) |  |
| Aarqa | *Enterobacter cloacae* (1)<br>*Klebsiella pneumoniae* (5)<br>*Hafnia alvei* (2)<br>*Serratia marcescens* (1)<br>*Serratia liquefaciens* (2)<br>*Salmonella choleraesuis* (1)<br>*Providencia rettgeri* (1)<br>*Serratia rubidaea* (1)<br>*Klebsiella oxytoca* (1)<br>*Pantoea spp.* (16) | *Klebsiella pneumoniae* (50) | *Pseudomonas luteola* (14)<br>*Pseudomonas fluorescens* (1)<br>*Pseudomonas putida* (1) |  | (11) |
| Damour | *Escherichia coli* (2) |  | *Pseudomonas luteola* (1) |  |  |
| Ostueene | *Enterobacter cloacae* (2)<br>*Hafnia alvei* (2) |  | *Pseudomonas luteola* (7)<br>*Pseudomonas putida* (1) |  | (1) |
| Zahrani | *Serratia liquefaciens* (3) |  |  |  |  |
| Kaleb |  |  | *Pseudomonas luteola* (2)<br>*Pseudomonas horizihabitans* (1) |  |  |
| Bared |  |  |  | *Pseudomonas putida* (3)<br>*Pseudomonas fluorescens* (1) |  |
| Beirut | *Enterobacter cancerogenus* (2)<br>*Serratia marcescens* (1)<br>*Klebsiella oxytoca* (1) | *Hafnia alvei* (10<sup>4</sup>) | *Pseudomonas luteola* (10<sup>4</sup>) | *Pseudomonas aeruginosa* (1) |  |
| Qadicha | *Serratia marcescens* (2)<br>*Hafnia alvei* (1)<br>*Cedecea lepagei* (1) |  | *Pseudomonas luteola* (1) |  |  |
| Awali |  | *Enterobacter amnigenus* (5)<br>*Escherichia coli* (2) |  | *Pseudomonas fluorescens* (3)<br>*Pseudomonas aeruginosa* (10) | (12) |

### Susceptibility profiles

We performed susceptibility testing for a panel of Enterobacterales (33) and *Pseudomonas* (39) strains. The results are shown in Table 2. We found 16 out of the 33 Enterobacterales and 6 out of the 39 *Pseudomonas* strains tested that expressed an ESBL phenotype detected with the synergy test.

**Table 2.**
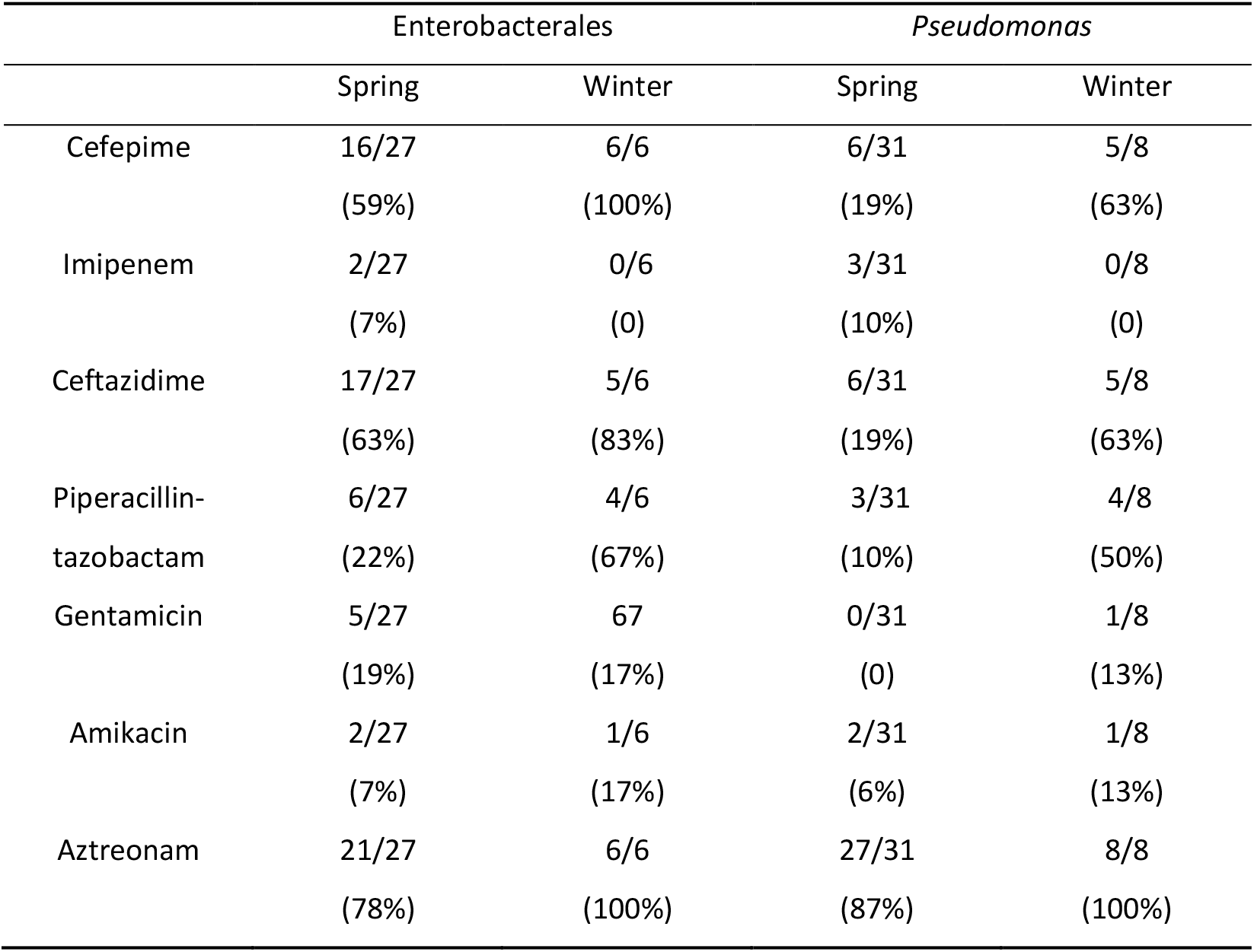
Resistance of the tested Enterobacterales and *Pseudomonas* species: numbers indicate the number of strains resistant out of the strains tested (%).

Resistance towards the tested antibiotics was broadly disseminated. All tested strains of *Pseudomonas* in spring were susceptible to gentamicin. In winter, we detected 2 *Pseudomonas aeruginosa* strains with an ESBL synergy test, 1 in Janoubi and 1 in Beirut. We detected 5 strains resistant to imipenem in spring, 3 ESBL positive *Pseudomonas luteola* strains (2 in Aarqa and 1 in Beirut), and 2 Enterobacterales strains (1 ESBL *Klebsiella oxytoca* in Aarqa and 1 *Serratia marcescens* in Qadicha). No resistant strains to imipenem were detected in winter.

### Resistome

Total DNA was extracted from triplicate river samples collected at the 10 estuaries of the major Lebanese rivers. Figure 1 depicts the abundance of each targeted gene normalized to the 16S rRNA encoding gene.

**Figure 1.**
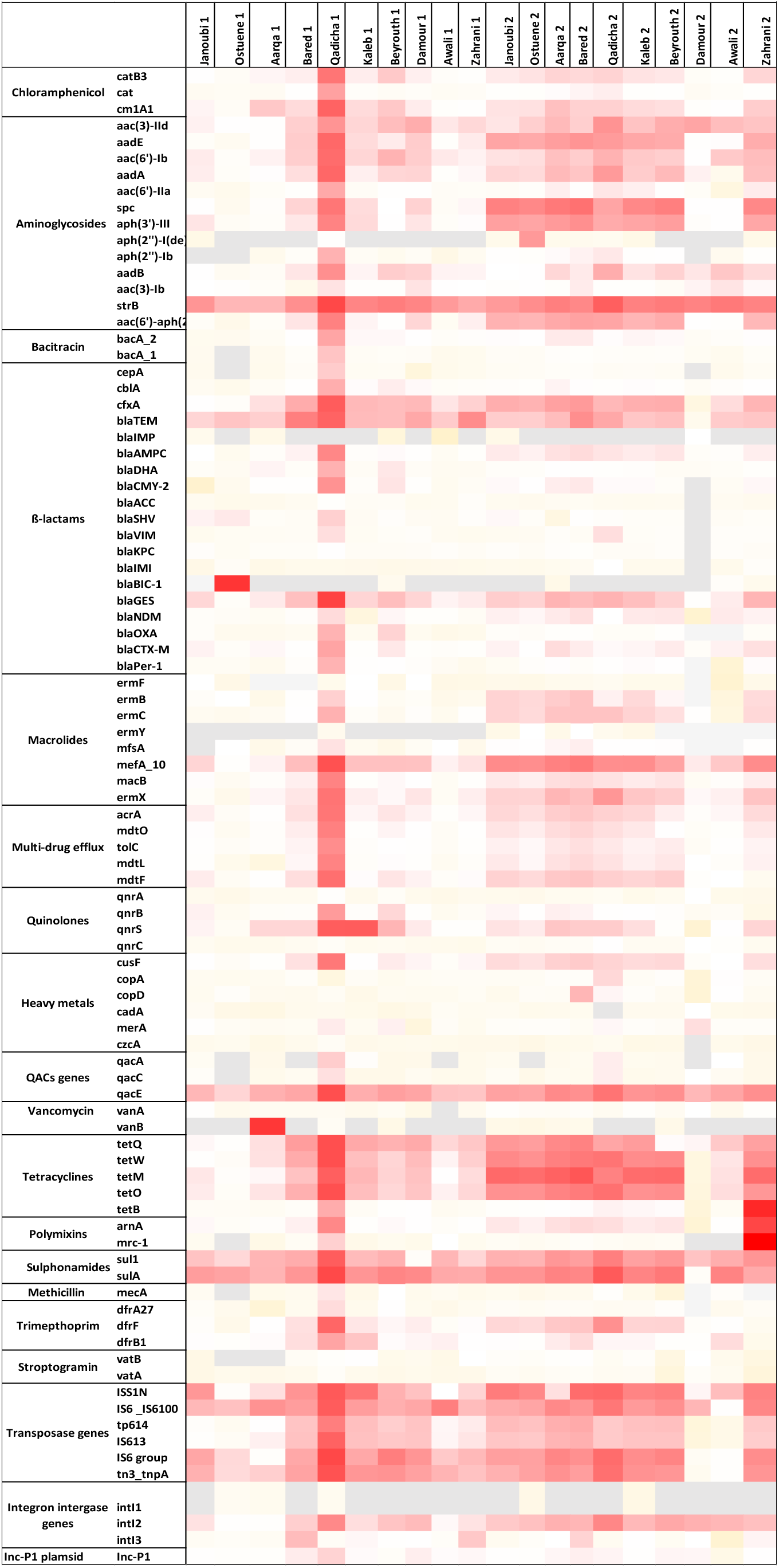
Heat map depicting the normalized abundances to the 16S rRNA gene of each targeted gene in spring (1) and in winter (2) in the 10 estuarine Lebanese rivers samples. Red: highest abundance, light yellow: lowest abundance, white medium, and grey: not detected

Overall, a higher normalized abundance of the targeted resistome was detected during winter, except for the Qadicha estuarine, in which high normalized abundance for ARGs was detected for all samples and seasons.

The IncP1 anthropogenic marker was detected in all samples at medium concentrations.

Interestingly, individual genes were detected in specific samples and sites to be highly abundant. For example, the highest normalized abundance in winter was detected for the *mcr-1* gene in the Zahrani river samples, followed by the *tetB* gene.

Noticeably, the *bla*_BIC-1_ and *tetM* genes were found in high normalized abundance in the Ostuene and Kaleb samples respectively. Also, during spring, the *vanB* gene was abundant in the estuarine samples from Aarqa, and the ESBL *bla*_GES_ gene was abundant in the Qadicha sample in spring.

The resistome signatures are different according to the season with a higher diversity in spring. A more conserved pattern is observed in winter, especially in the northern samples (Janoubi, Ostuene, Aarqa, and Bared). The resistome signatures from the middle section to the South of Lebanon, in addition to those from Qadicha, are more diverse.

The seasonal variation of the ARGs normalized abundance in the estuaries was assessed using the non-parametric Wilcoxon’s test. Figure 3 shows the estuaries where a significant increase of the mean normalized abundance was observed between spring and winter. The *p* values corresponding to all the significant variations are available in Supplementary Table 2.

Although we observed a significant increase for most of the estuaries, the mean normalized abundances of efflux pumps, tetracyclines, ß-lactams, heavy metal resistance genes in addition to the mobile genetic elements (MGEs) significantly decreased from spring to winter (*p* < 0.04) in the Qadicha and Damour estuaries. In Qadicha, the aminoglycoside and macrolide resistance genes decreased significantly (*p* < 0.04).

## Discussion

Previous studies underlined the contamination of Lebanese rivers with fecal bac-teria especially Escherichia coli and coliforms, reaching 70.4% of the rivers in the North and 60% in the Bekaa region, which was over the acceptable threshold they used ^30^.

Lebanon is located in the eastern Mediterranean area along a coastal length of 220 km ^29^ and is characterized by a short, cold, and wet winter season from January to March with annual rainfall ranging between 850 mm and 1800 mm/year, and by a dry summer ^31,32^. The surface waters in Lebanon are under increased pressure from anthropic activities, i.e. urban, industrial, and recreational. Agriculture in the coastal zone in Lebanon requires irrigation with surface and groundwater causing the depletion of water resources while increasing pollution according to the Food and Agriculture organization report of the year 2016 ^33^.

In our study, we showed that the relative abundance of the resistome varied according to the location. Figure 4 marks anthropogenic impact on the different geographic locations of Lebanon by indicating important urban zones, industrial zones, and activity, as well as zones with high population densities due to the influx of refugees. The Lebanese coastal belt from Tripoli to Tyre is the urbanized zone also witnessing agricultural activities. The highest count of bacteria, (Enterobacterales (*Hafnia alvei* (10^4^)) in winter and *Pseudomonas luteola* (10^4^) in spring) was observed in the estuary of the capital Beirut. The highest cumulated normalized abundance of ARGs was observed in the Qadicha estuary in spring which is consistent with a high industrial and urbanization influence in the area. The Qadicha river is a small river impacted by population growth, with industrial activities and a chronic default of wastewater treatment. It has been shown that the estuary of this river was highly contaminated with pollutants ^34^. The highest diversity of Enterobacterales was obtained from the Aarqa estuary in spring while a higher bacterial count was observed in winter (*Klebsiella pneumoniae* 50 UFC). The Aarqa estuary is located on the northern rural coast, inhabited by local and refugee communities with high agricultural activities. The elevated normalized abundance detected for the mobile colistin resistance gene *mcr-1* in the Zahrani estuary in winter could be linked to its closely located poultry production sites in Nabatyeh, where elevated numbers of *E. coli* carrying the *mcr-1* gene were isolated ^35^. The surface runoff from those broiler farms might impact and contaminate the Zahrani river with the *mcr-1* gene. Colistin is a polymixin antibiotic serving as the last-resort drug for carbapenemase-producing Enterobacterales - associated infections ^36^. The high abundance of the *mcr-1* gene is cause for concern as horizontal transfer to pathogens may occur under anthropogenic pressure ^37^.

We observed a high relative abundance of ARGs in the North (Figure 3) which corresponds to the area with a high density of refugees camps and with the highest number of inhabitants (Table 3). Lebanon hosts the largest number of refugees per capita worldwide. The Syrian crisis in the year 2011 has escalated the rivers’ water pollution, particularly on the northern coast following the migratory inflow of displaced persons searching for water points and settling to find refuge while infrastructure is lacking ^33^. This inflow has contributed to sustaining extreme hot spots of water stress in urban areas in informal settlements (the Northern coast or in the Bekaa region) while water networks and governance are already insufficient ^38^. In addition to the efforts from humanitarian organizations such as the UNHCR and the UNICEF to improve the sanitary of a fraction of the displaced people by the implementation of water and wastewater facilities, local initiatives to strengthen and/or rehabilitate existing infrastructure also exist and have to be continued to alleviate the vulnerability of displaced communities and to reduce the water stress in the country ^39,40^.

Generally, we could observe seasonal differences for the resistome and resistant bacterial isolates between spring and winter. River water levels in Lebanon increase during spring due to the rise of underground water and the snowmelt ^41^. This seasonal difference in water levels may contribute to the observed lower normalized abundances of the resistome in spring compared to the winter season (Figure 1). However, when analyzing the resistome signature, we observed a higher diversity of ARGs in spring (Figure 2). This could be explained by the fact that with the rise of temperature, people play, bathe, and do picnics along the beach and in estuaries, hence the observed diversity for the resistome in spring may reflect an impact due to recreational activities.

**Figure 2.**
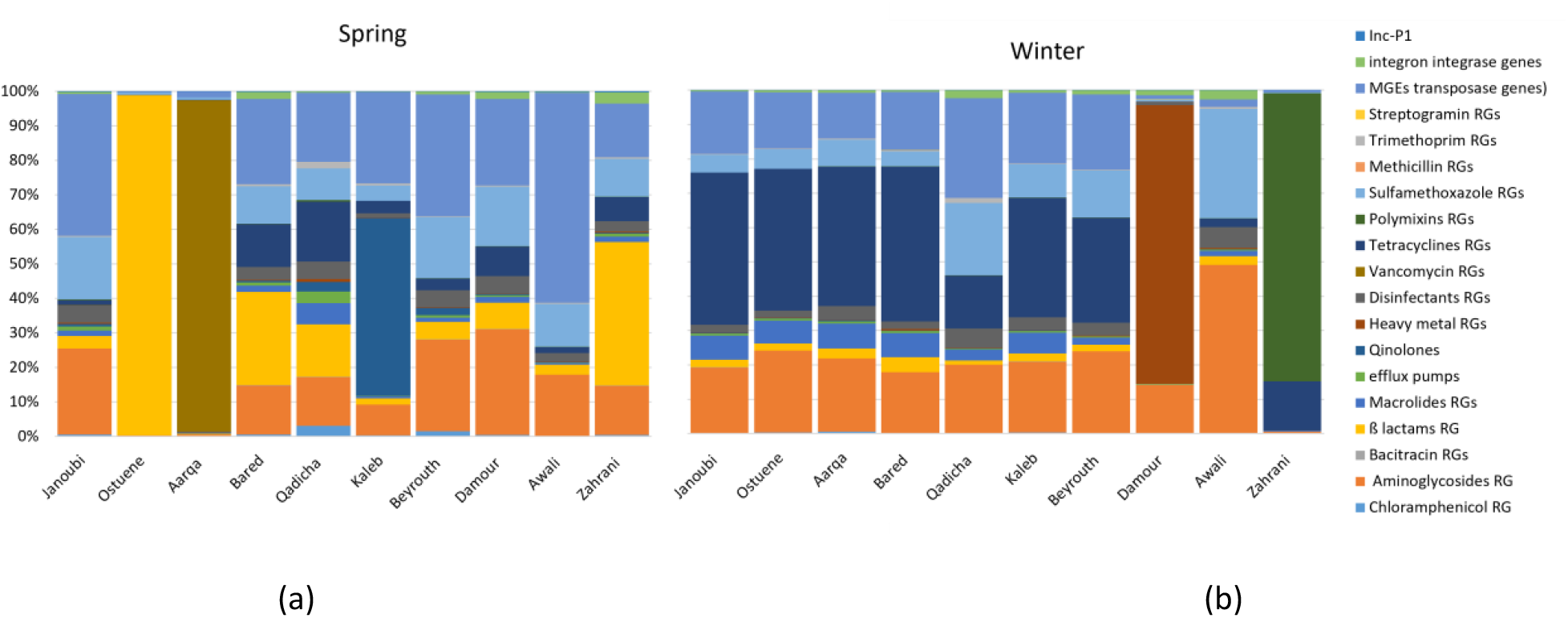
Proportional abundances of the variation of the ARGs families per river in (a) spring and in (b) winter in 10 estuarine Lebanese rivers samples

Most bacteria that were isolated from the rivers were highly resistant to antibiotics, especially to ß-lactams, while 16/33 Enterobacterales strains and 6/39 *Pseudomonas* strains expressed an ESBL phenotype. The resistome analysis showed a diversity of ESBL-encoding genes, including a high abundance of *bla*_IBC-1_ in the Ostuene river in North Lebanon. The *bla*_IBC-1_ gene was first characterized in 2002 in an *E. coli* strain in Greece, another country of the Mediterranean area ^42^. We also detected the ESBL gene *bla*_PER-1_ that was previously described in *P. aeruginosa* strains in other Middle East countries (Iran, Turkey, Tunisia)^43, 44, 45^, as well as the *bla*_CTX-M_ and *bla*_GES_ ESBL genes that are distributed worldwide in Enterobacterales and *Pseudomonas* strains. We also identified Enterobacterales and *Pseudomonas* strains resistant to imipenem. The resistome analysis led to the detection of a few *bla*_NDM_ and *bla*_OXA_ genes (including the carbapenemase OXA-48 encoding gene) in the rivers. Furthermore, it is known that some GES enzymes can also hydrolyze carbapenems ^46^. Altogether, the presence of ESBL and carbapenemase-producing pathogens and corresponding ARGs in the Lebanon estuaries raise concerns for public health, as in case of infection with these resistant bacteria, physicians have to rely on drug combinations and last-resort antibiotics ^47^.

Similar to our findings, multidrug-resistant Enterobacterales, namely *Enterobacter cloacae, Escherichia coli*, and *Klebsiella pneumoniae*, were isolated from the human-impacted Melayu river estuary in Malaysia with antibiotic resistance attaining 100% to antibiotics such as cefotaxime ^48^. Inversely, resistant Enterobacterales in the urbanized San Francisco Bay were absent from the near-shore sediments suggesting that urbanized estuaries may not constitute a major human exposure hazard when secondary and tertiary treatment operations and control measures for all wastewaters that drain into the studied environment are implemented ^49^.

The presence of MRSA in Aarqa, Ostuene, and Awali estuaries detected in spring should be closely monitored since studies have clearly shown that *Staphylococcus aureus* and MRSA persist in anthropized river samples presenting a potential source of the dissemination and transmission of resistant bacteria ^50,51,52^.

Currently, wastewater treatment plants (WWTP) in Lebanon have reported to mainly use secondary systems, although their status of operationality is undetermined. Two facilities in the Bekaa using a tertiary treatment system and 4 (3 in Mount Lebanon and 1 in the South) use a primary treatment system ^33^. The number of WWTP is proportional to the number of inhabitants, however, to prevent subsequent health effects from potential pathogens associated with poorly treated water ^53^, the number of WWTP using advanced technologies such as the tertiary treatment system that combines activated sludge with methods of reducing the bacterial load (disinfection, ultrafiltration) is scarce and should be increased.

Here, we identified the estuaries of Lebanon as putative host-spots for the dissemination of ARGs and resistant bacteria into the Mediterranean Sea. Recreational waters as rivers and beaches have gained increased attention as having a central role in the persistence, dissemination, and emergence of antibiotic resistance ^54^. Specific surveillance systems should be put in place that traces urban movements, contact with recreational waters, pollution levels, and WW treatment levels and management to trace the actual dissemination of antimicrobial resistance through recreational waters ^55^.

## Materials and Methods

Fifteen perennial Lebanese rivers spread through or at the extension of the Mount of Lebanon for a dozen of km eastward before discharging in small catchment areas leading to the Mediterranean Sea ^56^. The rivers spread over the territory making it a dense network of watercourses at a less than 10 km distance from each other and sharing similar basin characteristics ^56^. To evaluate the seasonal impact on the presence and abundance of ARBs and ARGs, sampling campaigns have been performed at two different periods.

Twelve estuaries or mouths were sampled in triplicate in sterile cups in April 2017 and January 2018 resulting in n=72 river water samples: 36 in spring and 36 in winter (3 samples of 60 ml each taken one after the other per river) (Figure 5). Weather conditions in terms of temperature with slight or no precipitation were similar along with the month duration for each of the two sampling campaigns. Samples were transported on ice directly to our Lebanese laboratory for further analysis within 2 hours. The Jawz river dries in coastal locations in the spring and therefore was not sampled for analysis. The exact coordinates registered for each sampling location as well as the temperatures are available in Supplementary Table 1.

**Figure 3.**
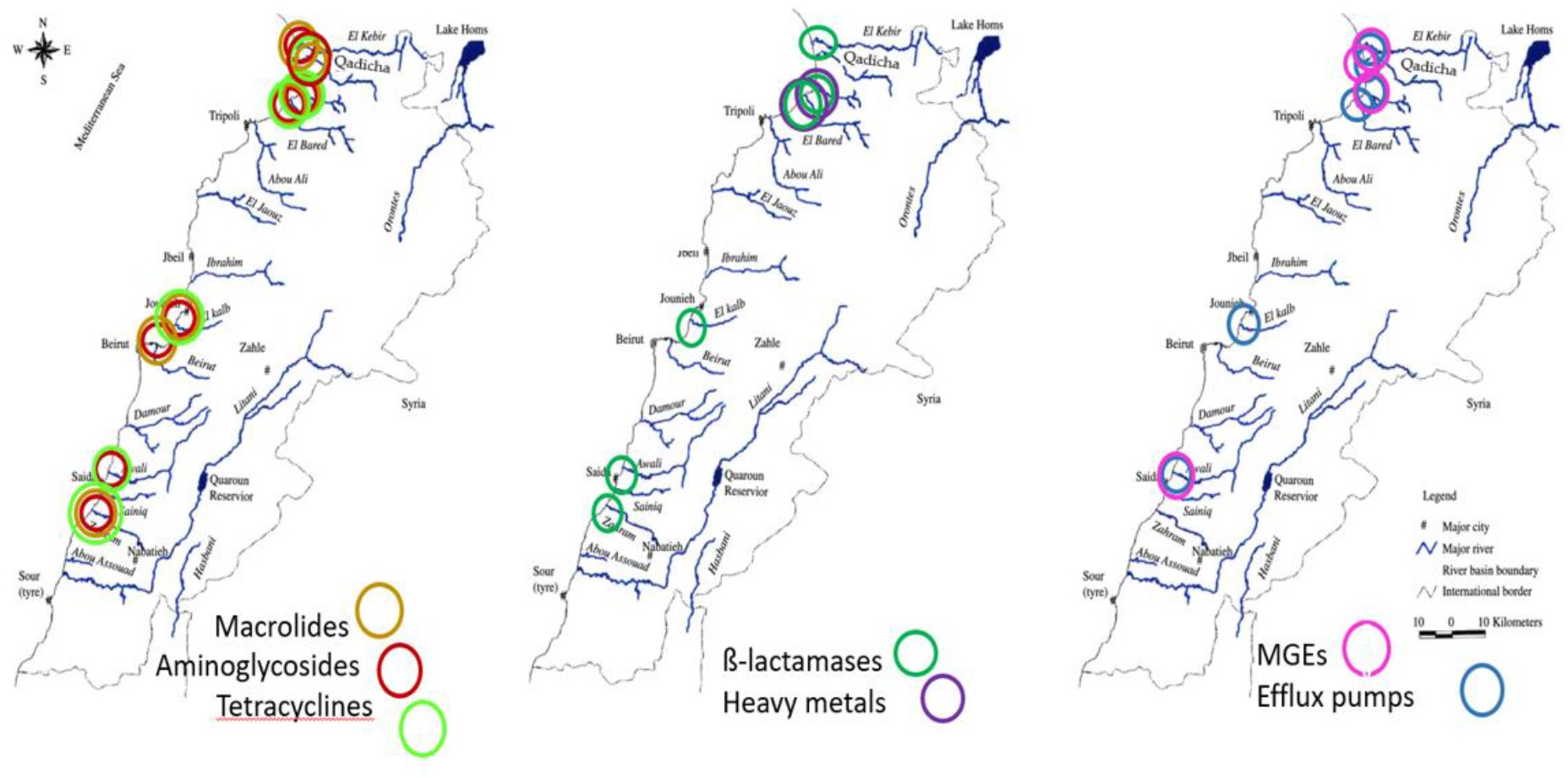
Graphical representation showing the estuaries where an increase in the mean normalized abundances of the ARGs families and MGEs from spring to winter was significant (p < 0.05).

**Figure 4.**
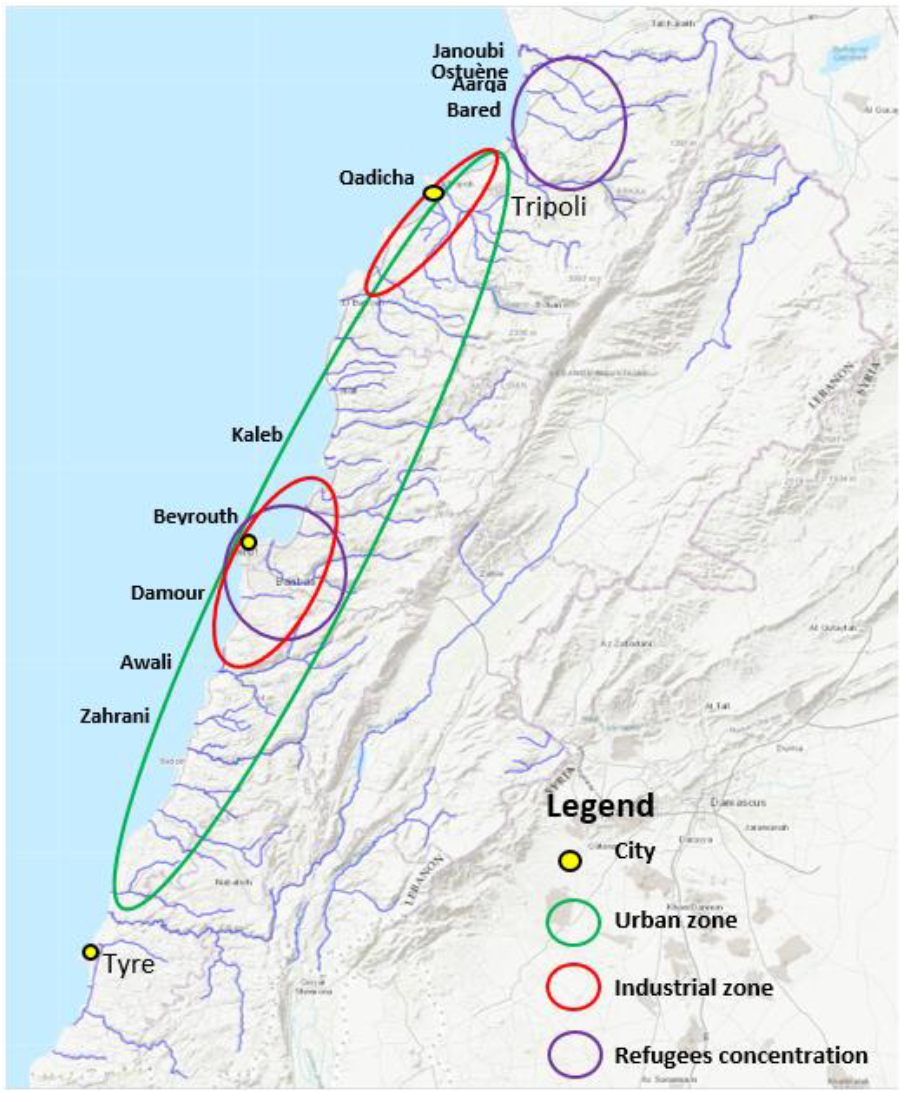
High anthropogenic impact locations. A map from “ Rivers, Lebanon, 2012. https://maps.princeton.edu/catalog/stanford-wn533df2039” representing the Lebanese rivers, modified to show locations on the Lebanese cost of the biggest industrial zones (Beirut and Tripoli), the highest urbanized area and location of refugees according to the UNHCR, UN-habitat 2014 and the UNRWA 2021 organizations.

**Figure 5.**
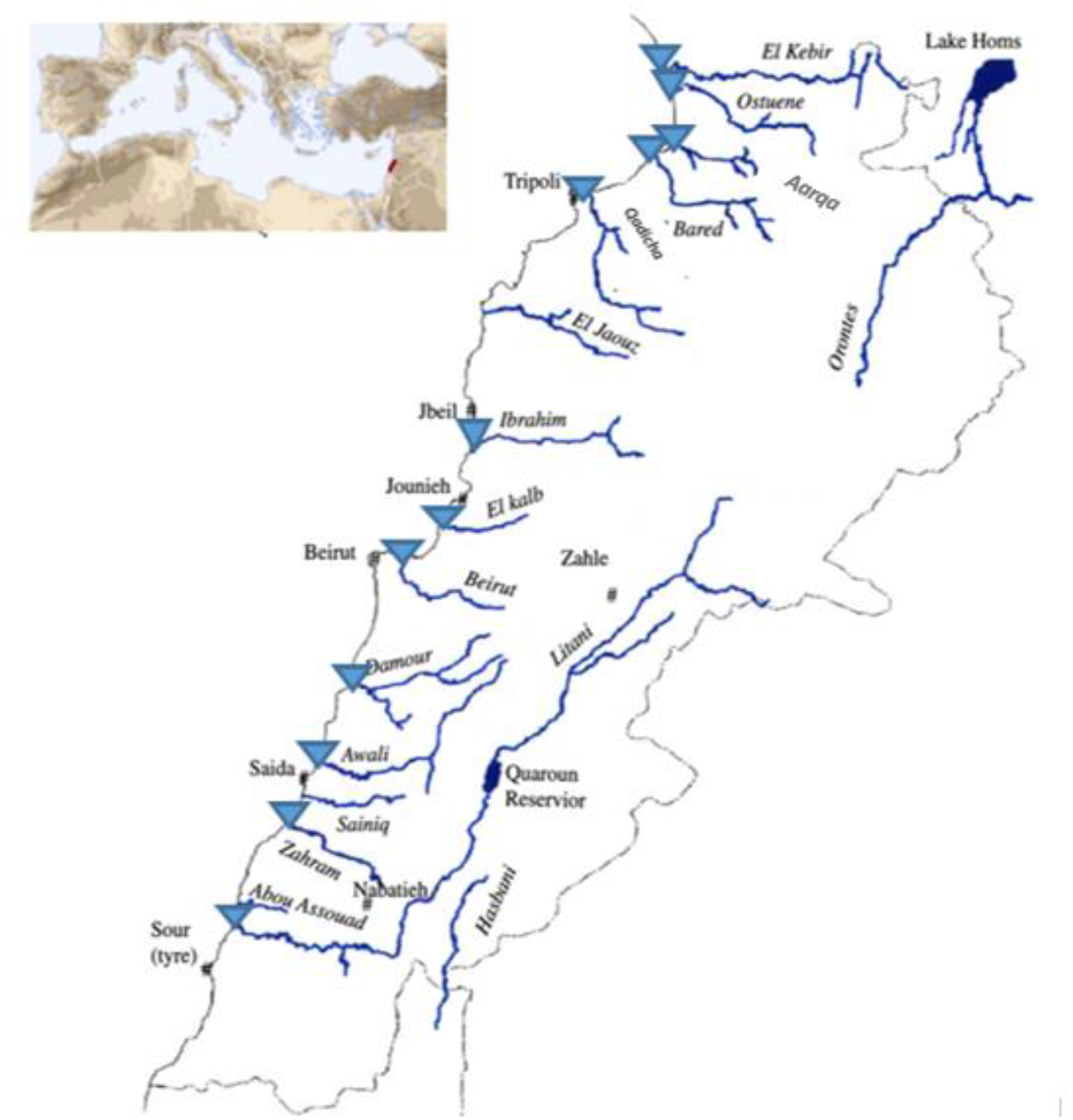
Sampling sites are represented as blue triangles on the map of the Lebanese rivers. Figure adapted from Houri & El Jeblawi, 2007

The water samples were cultured (V=1ml) using sterile rakes on Mac Conkey and Mannitol Salt agar selective of Gram-negative bacteria and *Staphylococcus spp*. respectively, with and without antibiotics. For Gram-negative bacteria, we used ceftriaxone (2 mg/L), cefepime (4 mg/L) or ertapenem (0.5 mg/L). For *Staphylococcus aureus*, we used oxacillin (4 mg/L). All media were incubated at 37 °C for 48 hours. For Gram-negative bacteria, species identification was performed with API® 20 NE (for *Pseudomonas*) or API® 20 E (for Enterobacterales) galleries (Biomérieux). For *S. aureus*, we used three phenotypic tests: catalase, DNAse, and coagulase.

The susceptibility testing was performed on Muller-Hinton agar according to the European Committee on Antimicrobial Susceptibility Testing (EUCAST) v8 guidelines. Antibiotic Bio-Rad® discs used were: amoxcicillin-clavulanic acid (20µg-10 µg), cefepim 30µg, ceftazidim (10µg), aztreonam (30µg), gentamicin (10µg), amikacin (30µg), piperacillin-tazobactam (30µg-6 µg) and imipenem (10 µg) for Gram-negative bacteria, and fusidic acid (10µg), cefoxitin (30µg), trimethoprim-sulfamethoxazole (1.25-23.75µg), gentamicin (10µg), and ciprofloxacin (5µg) for *S. aureus*.

For molecular analysis, the three water samples per river were filtered (total volume of water filtered = 180 ml), using a filtration ramp (Sartorius, Göttingen, Germany), on a sterile 47 mm diameter filter with a pore size of 0.45 µm (Sartorius, Göttingen, Germany). Microorganisms were recovered from filters and subject to DNA extraction for downstream analysis, using the DNeasy PowerWater® (Qiagen) adapted to water samples. All DNA samples were diluted or concentrated to a final concentration of 10 ng/µl for downstream qPCR and 16S rRNA analysis.

The Litani and Ibrahim rivers estuaries were excluded from the analysis as the respective water samples contained insufficient biomass even after an additional 2 L water volume sampling and DNA extraction.

We targeted 71 ARGs, 6 heavy metal resistance genes and 3 genes encoding resistance to quaternary ammonium compounds, 9 MGEs (transposases ISSW1, ISS1n, IS*6100*, IS*613*, IS*6* group, Tn*3*, ISCEc9, tp614), and integron integrase genes (*intI1, intI2*, and *intI3*) ^10^. The list of all the primers is published ^10^. The 16S rRNA encoding genes were targeted to allow normalizing the abundance of individual resistance genes.

The targeted genes are grouped according to their function ^10^ with, in addition, the added environmental resistance marker: incP1 ^57^.

High throughput real-time PCR was performed using the Biomark microfluidic system from Fluidigm in which every sample-gene combination is quantified using a 96.96 Dynamic Array(tm) IFCs (BMK-M-96.96, Fluidigm) as described previously ^58^.

A mean normalized abundance for each ARGs’ family and each estuary in spring and winter was calculated by dividing the cumulated abundance by the number of ARGs constituting the ARG family. The mean abundance of the ARGs families was then compared between spring and winter to assess a possible variation between the seasons. The estuaries where a significant variation was observed are listed in Supplementary Table 2.

## Conclusions

To date, this is the first study to provide an accurate assessment and comparison of the resistome in ten estuaries on the Lebanese coast. A combined approach using culture-based techniques and high throughput qPCR for the detection of ARBs and ARGs enabled identifying hotspots for antimicrobial resistance in the estuaries. This study highlighted the need to implement regular antibiotic resistance surveillance and improvement of waste management, in addition to the enforcement of regulations and guidelines stringency for sewage sludge or wastewater reuse as per the new EU 2020/741 legislation of the european parliament and of the council of 25 May 2020.

## Supporting information

Samples coordinates

Significant variations of ARGs families

## Supplementary Materials

**Table 1**: The exact coordinates of the sampling locations and the temperature recorded at the time of sampling, Table 2: Significant variations (p < 0.05) of ARGs families in the rivers between spring and winter.

## Author Contributions

W.H. designed the study, W.H. performed experiments, W.H., and E.B. performed data analysis, W.H., wrote the original draft, E.B. and M-C.P writing-review and editing with contribution of all other co-authors.

## Funding

This research was supported by The National Council for Scientific Research Lebanon CNRS-L, the Inserm and French Ministry of research, and the Research Council of the Saint Joseph University of Beirut.

## Competing interests

The authors declare no conflict of interest. The funders had no role in the design of the study; in the collection, analyses, or interpretation of data; in the writing of the manuscript, or in the decision to publish the results.

## Notes

### Competing Interest Statement

The authors have declared no competing interest.

