## Supplementary material for "Seasonal resistome diversity and dissemination of WHO priority antibiotic-resistant pathogens in Lebanese estuaries": Samples coordinates

Supplementray Table 1: The exact coordinates of the sampling locations and the temperature recorded at the time of sampling

| Fleuve | Coordonnées des points d'échantillonnage | Température en april 2017 en °C | Température en janvier 2018 en °C |
|---|---|---|---|
| Al Janoubi Al Kabir | 34° 38′ 04′′ N<br>35° 58′ 42′′ E | 19 | 13 |
| Ostuène | 34° 36′ 07′′ N<br>35° 59′ 24′′ E | 19 | 13 |
| Aarqa | 34° 32′ 58′′ N<br>35° 59′ 30′′ E | 19 | 13 |
| Al Bared | 34° 30′ 31′′ N<br>35° 57′ 42′′ E | 18 | 12 |
| Qadicha | 34° 26′ 48′′ N<br>35° 50′ 47′′ E | 20 | 12 |
| Ibrahim | 34° 03′ 54′′ N<br>35° 38′ 35′′ E | 20 | 13 |
| Al Kaleb | 33° 95′61′′ N<br>35° 58′ 80′′ E | 19 | 13 |
| Beyrouth | 33° 54′ 06′′ N<br>35° 32′ 18′′ E | 19 | 14 |
| Damour | 33° 42′ 17′′ N<br>35° 26′ 36′′ E | 18 | 15 |
| Al Awali | 33° 35′ 20′′ N<br>35° 23′ 12′′ E | 20 | 15 |
| Zahrani | 33° 29′ 33′′ N<br>35° 20′ 34′′ E | 18 | 14 |
| Litani | 33° 95′61′′ N<br>35° 59′80′′ E | 19 | 15 |
