## Supplementary material for "Seasonal resistome diversity and dissemination of WHO priority antibiotic-resistant pathogens in Lebanese estuaries": Significant variations of ARGs families

Supplementary Table 2: Significant variations (***p* < 0.05**) of ARGs families in the rivers between spring and winter

| | *Significant increase* ($p$ value) | *Significant decrease* ($p$ value) |
|---|---|---|
| Macrolides | Bared ($p$ 0.02) | Qadicha ($p$ 0.04) |
| | Ostuene ($p$ 0.02) | |
| | Beirut ($p$ 0.02) | |
| | Aarqa ($p$ 0.01) | |
| | Kaleb ($p$ 0.01) | |
| | Janoubi ($p$ 0.01) | |
| | Zahrani ($p$ 0.02) | |
| Aminoglycosides | Bared ($p$ *0)* | Qadicha ($p$ 0.04) |
| | Ostuene ($p$ *0)* | Damour ($p$ 0.04) |
| | Kaleb ($p$ *0)* | |
| | Aarqa ($p$ *0)* | |
| | Zahrani ($p$ *0)* | |
| Tetracyclines | Awali ($p$ 0.04) | Qadicha ($p$ 0.04) |
| | Bared ($p$ 0.04) | Damour ($p$ 0.04) |
| | Ostuene ($p$ 0.04) | |
| | Kaleb ($p$ 0.04) | |
| | Janoubi ($p$ 0.04) | |
| | Aarqa ($p$ 0.04) | |
| | Zahrani ($p$ 0.04) | |
| ß-lactamase | Awali ($p$ 0.01) | Qadicha ($p$ 0) |
| | Bared ($p$ 0.01) | Damour ($p$ 0) |
| | Kaleb ($p$ 0.01) | |
| | Aarqa ($p$ 0.01) | |
| | Janoubi ($p$ 0.01) | |
| | Zahrani ($p$ 0.01) | |
| Heavy metals | Aarqa ($p$ 0.03) | |
| | Bared ($p$ 0.03) | |
| MGEs | Aarqa ($p$ 0.01) | Qadicha ($p$ 0.02) |
| | Bared ($p$ 0.02) | Damour ($p$ 0.02) |
| | Janoubi ($p$ 0.02) | |
| | Zahrani ($p$ 0.04) | |
| Efflux | Awali ($p$ 0.04) | Qadicha ($p$ 0.04) |
| | Bared ($p$ 0.04) | Damour ($p$ 0.04) |
| | Ostuene ($p$ 0.04) | |
| | Kaleb ($p$ 0.04) | |
| | Janoubi ($p$ 0.04) | |
| | Aarqa ($p$ 0.04) | |
